# An Assessment of Skin Lesion Measurement Techniques for Use in Clinical Trials of Acute Bacterial Skin and Skin Structure Infections

**DOI:** 10.1101/312710

**Authors:** Michael W. Dunne, Anita Das, George H. Talbot

## Abstract

**Background:** Limited data are available to support a reproducible measurement technique that could be used to assess the response of a skin lesion associated with a bacterial infection to antibacterial therapy.

**Methods:** This multicenter, observational study enrolled patients with a major cutaneous abscess, a traumatic wound or surgical site infection, or a cellulitis. The primary objective was to characterize the intra- and inter-observer variability inherent in measuring the size of the erythema associated with the presenting skin infection. At least two observers made ruler measurements of the infection site with the length of the infection measured as the longest dimension of the erythematous area and width measured as the largest dimension perpendicular to the longest length. Intra- and inter-observer variability was determined by the intraclass correlation coefficient (ICC). Photographs, tracings, and thermal imaging were also performed.

**Results:** The intra*-*observer ICC (95% CI) for lesion area as measured by ruler was 0.999 (0.998, 0.999), suggesting only a very small amount of the variability was due to measurement error. The difference in mean lesion area measurements by the same observer was <1%. The inter-observer ICC (95% CI) for lesion area as measured by ruler was 0.990 (0.981, 0.995) suggesting that the results between observers were also highly reliable.

**Conclusions:** Measurement of infection area as defined by erythema and measured by ruler shows excellent intra- and inter-observer reliability and can be used in future clinical trials of acute bacterial skin infections.

## Introduction

Acute bacterial infections of the skin and skin structures (ABSSSI) are associated with both local and systemic signs of infection. Limited data are available regarding accurate and reproducible methods for measurement of the size of ABSSSI lesions when patients present for therapy. In the 2010 Draft Guidance for Industry issued by the Food and Drug Administration (FDA) for developing drugs for treatment of ABSSSI^1^, recently updated^2^, emphasis was placed on the measurement of the size of these infections to determine the relative efficacy of antibacterial therapies. The purpose of the current study was to compare various measurement techniques used to assess the size of a lesion caused by infection of the skin and skin structures.

## Methods

This study was a multicenter, observational study examining the signs and symptoms of ABSSSI that enrolled patients between November 23, 2010 and February 8, 2011. The primary objective was to characterize the intra- and inter-observer variability inherent in measuring the size of the erythema associated with the presenting skin infection (lesion) in patients with an ABSSSI. The primary outcome measures were the intra- and inter-observer variability of the measured area of the ABSSSI lesion at presentation, defined as length x width and as measured by ruler. Other potential measurement techniques were investigated, as detailed below.

### Patients

Patients between 18 - 85 years of age provided informed consent using an informed consent form approved by the investigative site’s Institutional Review Board. Each patient had an ABSSSI suspected or confirmed to be caused by one or more Gram-positive bacteria, including a major cutaneous abscess, a traumatic wound or surgical site infection, or a cellulitis, each of which exhibited associated erythema involving at least 75 cm^2^ of surface area. A major abscess was characterized as a collection of pus which required surgical incision and drainage and was defined by a margin of erythema that was >5 cm in all directions from the rim of induration or edema that defined the border of the abscess. A surgical site or traumatic wound infection was characterized by purulent drainage with surrounding erythema, edema and/or induration that occurred within 30 days after the trauma or surgery and exhibited a margin of erythema in at least one direction that was >5 cm from the edge of the infection. Cellulitis was defined as a diffuse skin infection characterized by spreading areas of erythema, edema and/or induration.

In addition to the requirement for visible erythema, all patients were required to have at least two of either purulent drainage/discharge, fluctuance, heat/localized warmth, tenderness to palpation, or swelling/induration, as well as at least one of either a white blood cell (WBC) count >12,000 cells/mm3, or a manual performed WBC differential count with >10% band forms, regardless of peripheral WBC count, or a fever (body temperature ≥38°C/100.4°F); at least 25% of patients enrolled were to have a fever at baseline.

Patients were excluded if they were in shock, had an infection due to a Gram-negative bacterium, a venous catheter entry site infection, a diabetic foot ulceration, a decubitus ulcer, an infected device, or a full or partial thickness burn; or had a superficial/simple cellulitis, an impetiginous lesion, a furuncle, or a simple abscess that only required surgical drainage for cure.

### Infection site assessments

Infection site assessment included the following: presence or absence of purulence/drainage, erythema, heat/localized warmth, tenderness to palpation, fluctuance and swelling/induration.

#### Ruler measurement

At least two, but no more than three, observers with prior experience in clinical trials of ABSSSI made two assessments of the infection site. The first observer measured the erythema associated with the infection twice, with the second measurement separated from the first by 15 minutes. Each observer measured the infection erythema independently of the others, and was not present when other observers made their recordings. The erythema length and width were measured by ruler with grading by millimeters. The length of the infection was measured as the longest dimension of the erythematous area. Width was measured as the largest dimension perpendicular to the longest length. A standard paper tape measure was supplied to each site with gradations in millimeters. The primary assessment was based on the area defined by the length of the erythematous lesion multiplied by its width.

#### Camera measurement

Photographic images taken with an ARANZ Medical Silhouette Camera^™^ (Aranz, Ltd., Christchurch, New Zealand) were obtained that included an assessment of the margins of the infection using laser guided sightings such that a digital measurement of the lesion surface area could be obtained.

#### Tracings

Two-ply transparent acetate tracing material (Canfield Scientific Inc., Fairfield New Jersey, USA) was applied to the infection site and the margins of the erythematous area traced onto the transparency with a black marker. Two additional markings were made identifying the head-to-toe axis of the lesion. The transparencies were then read by planimetry to calculate the lesion area.

#### Thermal imaging

Thermal imaging was performed at selected investigative sites using an ICI thermal camera, model 7640 (ICI, Beaumont, Texas).

### Statistical Methodology

Descriptive statistics (mean, standard deviation, median, minimum and maximum) of the length, width, and area (length x width) based on a ruler measurement were generated for the first and second observers (and, if available, the third). The intraclass correlation coefficient (ICC) was determined for the two measurements completed by the first observer to assess the intra-observer variability and was determined for the measurements taken by the different observers to assess the inter-observer variability. The ICC was calculated by fitting an analysis of variance (ANOVA) model. Ninety-five percent confidence intervals (CI) for the ICC were constructed. The coefficient of variation (COV) for each measurement (length, width and area) is presented with the 95% CI. Comparison was also made of the erythema size as measured by the digital camera with that of the transparency by calculating the ICC (and 95% CI) and with a paired t-test.

Interpretation of the ICC was based on the proposal by Landis and Koch^3^, in which an ICC=0 is defined as having no reliability, 0<ICC≤0.2 as slight reliability, 0.2<ICC≤0.4 as fair reliability, 0.4<ICC≤0.6 as moderate reliability, 0.6<ICC≤0.8 as substantial reliability, and 0.8<ICC≤1.0 as almost perfect reliability.

Approximately 40 adult patients were to be enrolled with the sample size calculation based on the width of the 95% CI for the ICC using the method outlined by Bonnet^4^. If two observers provide lesion measurements, with a sample size of 40 patients and an estimated ICC of 0.90, the 95% CI will have a lower bound of 0.78.

## Results

Thirty-nine patients who met the inclusion and exclusion criteria were enrolled. The mean age was 43.7 years and 69.2% were Caucasian (Table 1). Patients had recent trauma (30.8%), diabetes mellitus (30.8%) and/or intravenous drug abuse (25.9%). Among the systemic signs of infection, 7 (17.9%) were febrile and 37 (94.7%) had a WBC count greater than 12,000 cells/mm^3^. The most common site of infection was the leg (35.9%). Fifteen patients had cellulitis, 19 a major cutaneous abscess, and 5 a traumatic wound infection. All patients had an infection site that was warm to the touch and evidenced erythema and tenderness; all but one demonstrated swelling or induration (Table 2).

**Table 1.** Baseline Demographics, Systemic Signs and Symptoms

|  | All Patients<br>N=39 |
| --- | --- |
| Age (years) |  |
| Mean (SD) | 43.7 (14.75) |
| Median (Min, Max) | 42.0 (20, 79) |
| Male | 25 ( 64.1) |
| Race, n (%) |  |
| White | 27 ( 69.2) |
| Black, African American, or African Heritage | 8 ( 20.5) |
| Other Native Hawaiian or other Pacific Islander | 4 ( 10.3) |
| Ethnicity, n (%) |  |
| Hispanic or Latino | 13 ( 33.3) |
| Not Hispanic or Latino | 26 ( 66.7) |
| Past Medical History, n (%) |  |
| Recent trauma | 12 ( 30.8) |
| Diabetes Mellitus | 12 ( 30.8) |
| Current or recent intravenous drug use | 10 ( 25.6) |
| Current or recent alcohol abuse | 4 ( 10.3) |
| Prior skin infection | 4 ( 10.3) |
| Peripheral vascular disease | 1 ( 2.6) |
| Temperature (°C), mean (SD) | 37.2 (0.84) |
| Febrile, n (%) | 7 (17.9) |
| WBC $\geq 12,000$ (cells/mm<sup>3</sup>), n (%) | 37 (94.5) |
| Immature Neutrophils $\geq 10$ (%), n (%) | 0 |
| C-reactive Protein (mg/dL), median (range)<sup>1</sup> | 9.8 (0.5, 288.5) |
| Assessment of Pain, median ( range) | 7.0 (1, 10) |
| Lymphadenopathy proximal to ABSSSI, n (%) | 27 ( 69.2) |
| Painful lymphadenopathy proximal to ABSSSI, n (%) | 22 ( 56.4) |

**Table 2.** Local signs of infection [n (%)]

| Sign/Symptom | Abscess<br>N=19 | Surgical Site/<br>Traumatic Wound<br>N=5 | Cellulitis<br>N=15 | All Patients<br>N=39 |
| --- | --- | --- | --- | --- |
| Erythema | 19 (100) | 5 (100) | 15 (100) | 39 (100) |
| Heat/Localized Warmth | 19 (100) | 5 (100) | 15 (100) | 39 (100) |
| Tenderness to Palpation | 19 (100) | 5 (100) | 15 (100) | 39 (100) |
| Swelling | 13 (68.4) | 2 (40.0) | 9 (60.0) | 24 (61.5) |
| Induration | 19 (100.0) | 5 (100.0) | 14 (93.3) | 38 (97.4) |
| Fluctuance | 9 (47.4) | 2 (40.0) | 1 (6.7) | 12 (30.8) |
| Discharge/ Drainage | 16 (84.2) | 5 (100.0) | 2 (13.3) | 23 (59.0) |

The mean infection size (based on measurement 1 by the first observer) was approximately 525 cm^2^ with a median size of approximately 350 cm^2^, reflecting a wide variability in the area of erythema for different patients (Table 3). Across the 39 patients enrolled, the *intra-*observer ICC (95% CI) for lesion area as measured by ruler was 0.999 (0.998, 0.999), suggesting extremely high reliability (i.e., only a very small amount of the variability was due to measurement error). The ICC (95% CI) for both length [0.996 (0.993, 0.998] and width [0.996 (0.993, 0.998)] was similar to that of lesion area. The difference in mean lesion area measurements by the same observer was <1% (524.988 cm^2^ vs. 526.124 cm^2^).

**Table 3.**
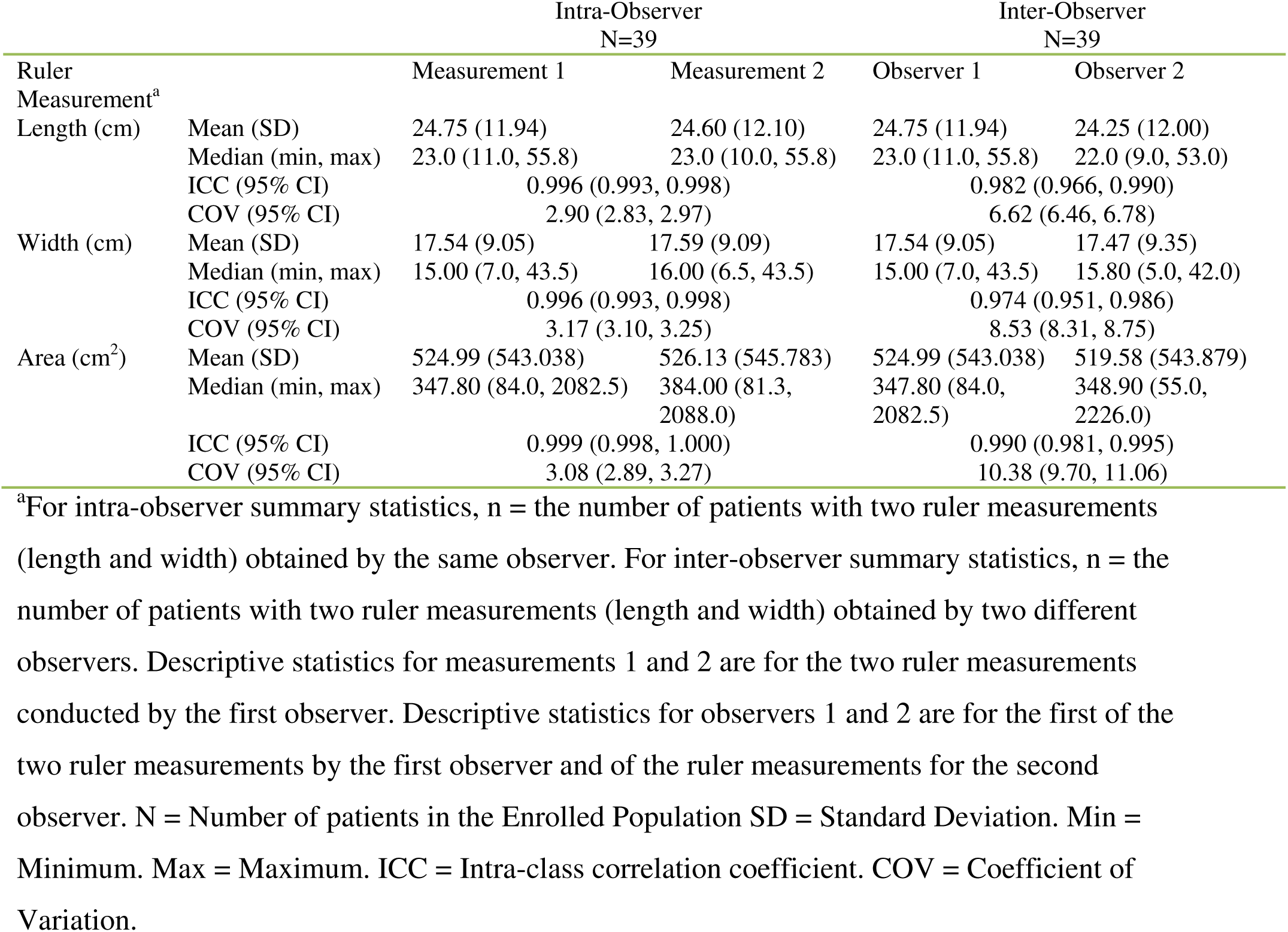
Intra-Observer and Inter-Observer Variability in Ruler Measurements of ABSSSI Lesion

|  |  | Intra-Observer<br>N=39 |  | Inter-Observer<br>N=39 |  |
| --- | --- | --- | --- | --- | --- |
| Ruler Measurement<sup>a</sup> |  | Measurement 1 | Measurement 2 | Observer 1 | Observer 2 |
| Length (cm) | Mean (SD) | 24.75 (11.94) | 24.60 (12.10) | 24.75 (11.94) | 24.25 (12.00) |
|  | Median (min, max) | 23.0 (11.0, 55.8) | 23.0 (10.0, 55.8) | 23.0 (11.0, 55.8) | 22.0 (9.0, 53.0) |
|  | ICC (95% CI) | 0.996 (0.993, 0.998) |  | 0.982 (0.966, 0.990) |  |
|  | COV (95% CI) | 2.90 (2.83, 2.97) |  | 6.62 (6.46, 6.78) |  |
| Width (cm) | Mean (SD) | 17.54 (9.05) | 17.59 (9.09) | 17.54 (9.05) | 17.47 (9.35) |
|  | Median (min, max) | 15.00 (7.0, 43.5) | 16.00 (6.5, 43.5) | 15.00 (7.0, 43.5) | 15.80 (5.0, 42.0) |
|  | ICC (95% CI) | 0.996 (0.993, 0.998) |  | 0.974 (0.951, 0.986) |  |
|  | COV (95% CI) | 3.17 (3.10, 3.25) |  | 8.53 (8.31, 8.75) |  |
| Area (cm<sup>2</sup>) | Mean (SD) | 524.99 (543.038) | 526.13 (545.783) | 524.99 (543.038) | 519.58 (543.879) |
|  | Median (min, max) | 347.80 (84.0, 2082.5) | 384.00 (81.3, 2088.0) | 347.80 (84.0, 2082.5) | 348.90 (55.0, 2226.0) |
|  | ICC (95% CI) | 0.999 (0.998, 1.000) |  | 0.990 (0.981, 0.995) |  |
|  | COV (95% CI) | 3.08 (2.89, 3.27) |  | 10.38 (9.70, 11.06) |  |

<sup>a</sup>For intra-observer summary statistics, n = the number of patients with two ruler measurements (length and width) obtained by the same observer. For inter-observer summary statistics, n = the number of patients with two ruler measurements (length and width) obtained by two different observers. Descriptive statistics for measurements 1 and 2 are for the two ruler measurements conducted by the first observer. Descriptive statistics for observers 1 and 2 are for the first of the two ruler measurements by the first observer and of the ruler measurements for the second observer. N = Number of patients in the Enrolled Population SD = Standard Deviation. Min = Minimum. Max = Maximum. ICC = Intra-class correlation coefficient. COV = Coefficient of Variation.

The *inter*-observer ICC (95% CI) for lesion area as measured by ruler was 0.990 (0.981, 0.995) suggesting that the results between observers were also highly reliable (Figure 1). The ICC (95% CI) for both length [0.982 (0.966, 0.990)] and width [0.974 (0.951, 0.986)] was similar to that of lesion area. The difference in mean lesion area measurements between two observers was 1% (524.988 cm^2^ vs. 519.584 cm^2^).

**Figure 1.**
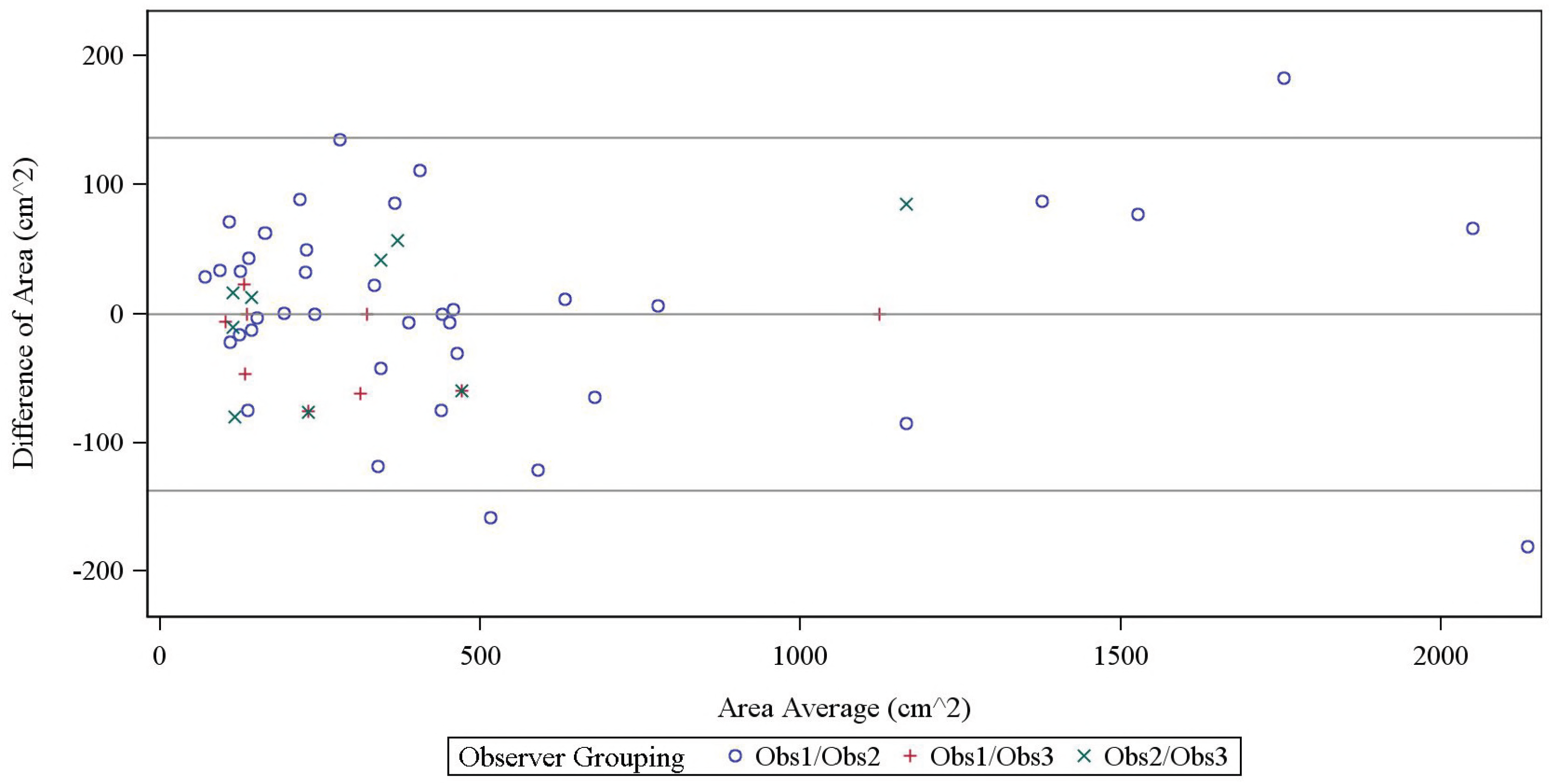
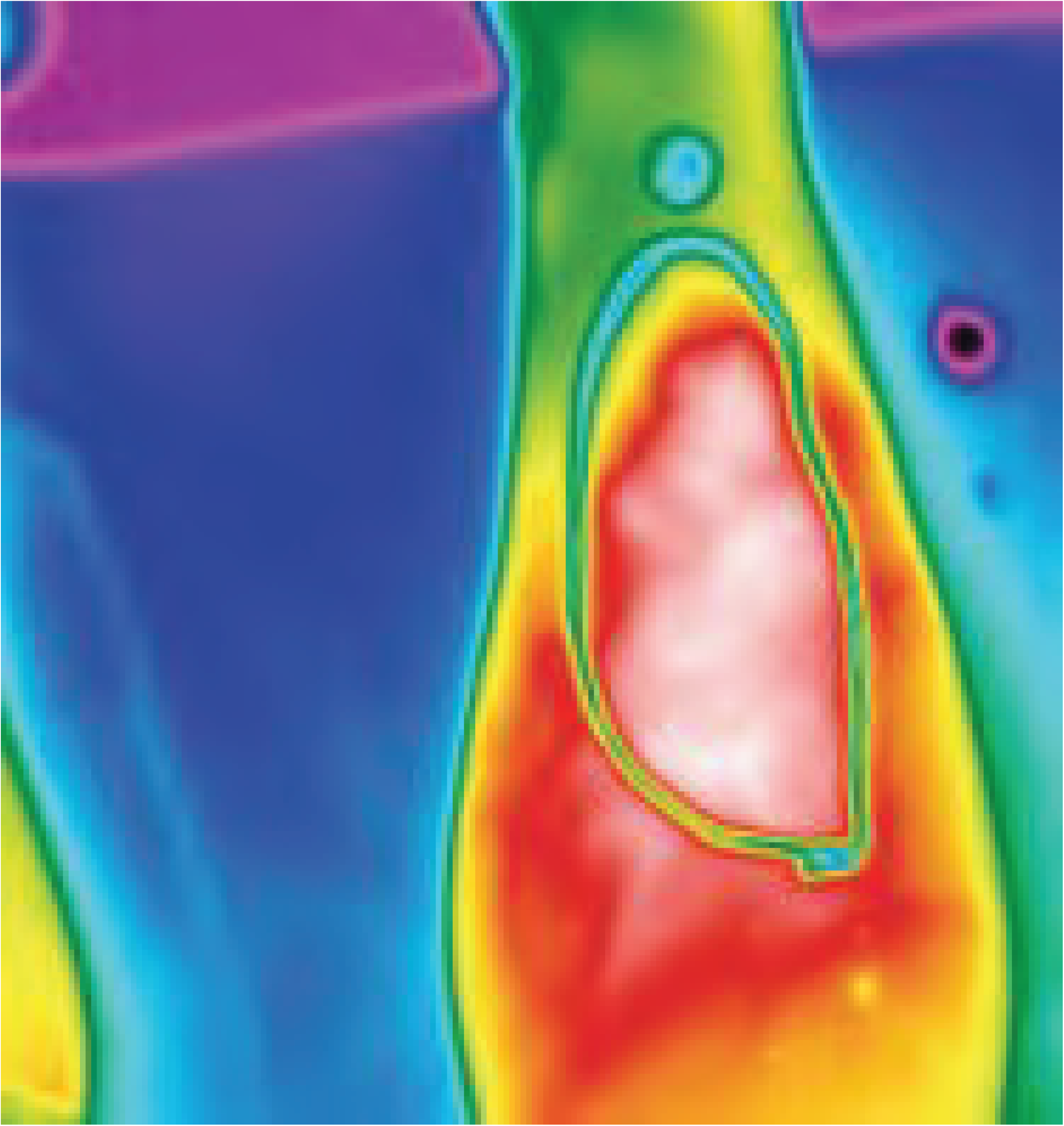
Bland-Altman Plot of Inter-Observer Ruler Area Measurements

The mean area of the infection as measured by either the digital camera or tracing was smaller than the infection size as measured by the ruler (303 cm^2^, 360 cm^2^, and 525 cm^2^, respectively). The ICC for the comparison of infection size as measured by the digital camera compared to the tracings was 0.880 (0.773, 0.939), p=0.04 (Table 4).

**Table 4.** Comparison of Lesion Size as Measured by Digital Image and Transparency

| Measurement |  | Patients | Paired Difference <sup>a</sup> |
| --- | --- | --- | --- |
| Digital Image (cm<sup>2</sup>) | n | 32 | 0 |
|  | Mean (SD) | 302.56 (295.50) | N/A |
|  | Median (min, max) | 199.10 (43.7, 1195.1) | N/A |
| Transparency (cm<sup>2</sup>) | n | 32 | 32 |
|  | Mean (SD) | 360.17 (360.74) | -57.61 (153.63) |
|  | Median (min, max) | 249.30 (60.5, 1362.5) | -21.35 (-552.8, 162.3) |
| Comparison | p-value <sup>b</sup> |  | 0.0420 |
|  | ICC (95 % CI) |  | 0.880 (0.773, 0.939) |

<sup>a</sup>Paired difference is calculated as digital imaging measurement minus transparent dressing measurement within each patient. <sup>b</sup>Paired t-test.

Thermal imaging provided additional qualitative information related to the heat signature associated with a specific lesion. Of interest, the area of warmth frequently exceeded the observed area of erythema.

## Discussion

The management of patients with cellulitis has typically relied upon an assessment of erythema size in order to make decisions related to efficacy of therapy^5^. Clinicians will frequently outline the lesion with an indelible marker to track for progression or regression of the infection over time. While lesion size measurements have been central to clinical care, very little information is available in the literature to support a reliable and reproducible measurement technique that could be used in registrational clinical trials.

The results from this study confirm that measurement with a paper tape of the erythema associated with a skin infection, specifically the longest length multiplied by the widest width perpendicular to that longest length, provides a reliable measurement of infection size both when assessed by the same observer over two occasions or across observers. These data, generated from an inexpensive and simple procedure, validate an approach to patient care that has been in practice for generations.

Roughly 25% of paired measurements differed by more than 20% across observers, concentrated mostly in lesions between 100 cm^2^ and 250 cm^2^. As a result, interpretation of a treatment response, either absolute or relative to another drug, should be made with some caution when the difference in lesion area between two measurements is <50 cm^2^. Typically over 90% of patients will have achieved a reduction of 90% in their lesion size by 72 hours post the initiation of treatment so this degree of variability in the measurement of lesion size is unlikely to be of clinical significance.

The availability of a simple and reproducible measurement technique is also of use to studies attempting to demonstrate efficacy of a new agent relative to an established one^6^. The 2013 FDA Final Guidance for Treatment of ABSSSI now relies on an assessment of the reduction in erythema size of ≥20% at 48-72 hours post-baseline as the primary endpoint in registration studies^2^. The data presented here provide support for a measurement technique to be used in such trials.

It is obvious that the estimation of the area of a skin infection by a length by width calculation will always overestimate the true size of that lesion except in the rare case when the erythema is in the shape of a perfect rectangle. For the purpose of estimating a response to treatment, the actual starting size of the infection is less important than the change in the size over time, as has been established in numerous trials over the last few years^7, 8, 9, 10^. Measurements obtained by planimetry of the tracing or with the digital camera, however, will provide an accurate estimation of the actual size of the lesion. The overestimation of actual lesion size based on a ruler measurement relative to these techniques is 30- 40%, which is approximately the relative difference of the size of a rectangle and an oval contained within its borders.

The purpose of this study was to provide as assessment of the accuracy of lesion measurement by means of a ruler. Other techniques were included as part of the trial, and provide some additional points of reference. Measurements by tracings or cameras, while potentially more reflective of the actual lesion size, are also more problematic. Because the area of the infection does not lend itself to being contained within one frame, multiple pictures must be carefully ‘stitched’ together. While software programs can make these calculations, careful preparation of the outline of these various sections must be performed on the patient’s skin, something that is not possible for every lesion in every patient. Tracings are simpler; however, again, certain lesions do not conform to the acetate because of their large size or the location of the infection and prevent an accurate representation. While the ruler is simplest, all three techniques provide complimentary utility and could be considered for use in the conduct of clinical trials for ABSSSI.

In conclusion, measurement of the size of an ABSSSI can be performed reliably by use of a ruler with a high degree of reliability within and among observers.

## Supplementary Data

Supplementary materials are available at *Antimicrobial Agents and Chemotherapy* online.

## Acknowledgements

The authors would like to acknowledge the contribution of the participating investigators: Purvi Mehra (Chula Vista, CA), Sinikka Green (La Mesa, CA), Wade Sears (Las Vegas, NV), Robert Cockrell (Fountain Valley, CA) and Paul Manos (Oceanside, CA).

## References

1. US Food and Drug Administration, Center for Drug Evaluation. (2010, August). “Guidance for Industry: Acute Bacterial Skin and Skin Structure Infections: Developing Drugs for Treatment.” Available at http://www.fda.gov/downloads/Drugs/GuidanceComplianceRegulatoryInformation/Guidances/ucm071185.pdf; replaced on this website by reference 2, below.

2. US Food and Drug Administration, Center for Drug Evaluation. (2013, October). “Guidance for Industry: Acute Bacterial Skin and Skin Structure Infections: Developing Drugs for Treatment.” Available at http://www.fda.gov/downloads/Drugs/GuidanceComplianceRegulatoryInformation/Guidances/ucm071185.pdf)

3. Landis J, Koch G. The measurement of observer agreement for categorical data. Biometrics 1977; 33:159–174.

4. Bonnet DG. Sample size requirements for estimating intraclass correlations with desired precision. Statist Med 2002; 21:1331–1335.

5. Snodgrass WR, Anderson T. Sulphanilamide in the treatment of erysipelas. Br Med J 1937; 2:1156–1159.

6. Talbot GH, Powers JH, Fleming TR et al. Progress on developing endpoints for registrational clinical trials of community-acquired bacterial pneumonia and acute bacterial skin and skin structure infections: Update from the Biomarkers Consortium of the Foundation for the National Institutes of Health. Clin Infect Dis 2012; 55:1114–1121.

7. Boucher HW, Wilcox M, Talbot GH, Puttagunta S, Das AF, and Dunne MW. Once-Weekly Dalbavancin versus Daily Conventional Therapy for Skin Infection. N Engl J Med 2014; 23:2169–2178.

8. Corey GR, Kabler H, Mehra P, Gupta S, Overcash JS, Porwal A, Giordano P, Lucasti C, Perez A M.D., Good S, Jiang H, Moeck G, and O’Riordan W, for the SOLO I Investigators. Single-Dose Oritavancin in the Treatment of Acute Bacterial Skin Infections. N Engl J Med 2014; 370:2180–2190.

9. Prokacimer PP, Anda CD, Fang E, Mehra P, Das A. Tedizolid Phosphate vs Linezolid for Treatment of Acute Bacterial Skin and Skin Structure Infections The ESTABLISH-1 Randomized Trial. JAMA 2013; 309: 559–569.

10. Dunne MW, Puttagunta S, Giordano P, Krievins D, Zelasky M, Baldassarre J. A Randomized Clinical Trial of Single-Dose Versus Weekly Dalbavancin for Treatment of Acute Bacterial Skin and Skin Structure Infection. Clin Infect Dis 2016; 62:545–551.

